# Calcium-driven regulation of voltage-sensing domains in BK channels

**DOI:** 10.1101/520429

**Authors:** Yenisleidy Lorenzo-Ceballos, Willy Carrasquel-Ursulaez, Karen Castillo, Osvaldo Alvarez, Ramon Latorre

## Abstract

Allosteric interplays between voltage-sensing domains (VSD), Ca^2+^-binding sites, and the pore domain govern the Ca^2+^- and voltage-activated K^+^ (BK) channel opening. However, the functional relevance of the Ca^2+^- and voltage-sensing mechanisms crosstalk on BK channel gating is still debated. We examined the energetic interaction between Ca^2+^ binding and VSD activation measuring and analyzing the effects of internal Ca^2+^ on BK channels gating currents. Our results indicate that the Ca^2+^ sensors occupancy has a strong impact on the VSD activation through a coordinated interaction mechanism in which Ca^2+^ binding to a single α-subunit affects all VSDs equally. Moreover, the two distinct high-affinity Ca^2+^-binding sites contained in the C-terminus domains, RCK1 and RCK2, appear to contribute equally to decrease the free energy necessary to activate the VSD. We conclude that voltage-dependent gating and pore opening in BK channels is modulated to a great extent by the interaction between Ca^2+^ sensors and VSDs.

## Introduction

Diverse cellular events involve calcium ions as a primary mediator in the signal transduction pathways triggering, among other signaling processes, Ca^2+^-activated conductances. Since the BK channels are regulated by cytosolic Ca^2+^ and depolarizing voltages (Marty, 1981; Pallotta et al., 1981; Latorre et al., 1982), they are integrators of physiological stimuli including intracellular Ca^2+^ elevation and membrane excitability. BK channels are modular proteins where each module accomplishes a specific channel function. Thus, different modules harbor voltage and Ca^2+^ sensors that communicate with the channel gate allosterically (Cox et al., 1997; Horrigan and Aldrich, 1999, 2002; Horrigan et al., 1999; Rothberg and Magleby, 1999, 2000; Cui and Aldrich, 2000). Functional BK channels are formed by homotetramers of α-subunits (Shen et al., 1994) each comprising a transmembrane voltage-sensing domain (VSD) and an intracellular Ca^2+^-sensing C-terminal domain (CTD) that can independently modulate the ion conduction gate in the pore domain (PD) (Latorre et al., 2017). The CTDs consist of two non-identical regulators of the conductance of K^+^ domains (RCK1 and RCK2) arranged into a ring-like tetrameric structure dubbed the gating ring (Wu et al., 2010; Yuan et al., 2010, 2012; Hite et al., 2017; Tao et al., 2017). Each RCK domain contains distinct ligand-binding sites capable of detecting Ca^2+^ in the micromolar range (Schreiber and Salkoff, 1997; Bao et al., 2002; Xia et al., 2002).

In the absence of Ca^2+^, the activation of VSD decreases the free energy necessary to fully open the BK channels in an allosteric fashion (Horrigan and Aldrich, 1999; Horrigan et al., 1999). Under these experimental conditions, very positive membrane potentials are required to drive all voltage sensors to its activated conformation (Cui et al., 1997; Stefani et al., 1997; Horrigan et al., 1999; Contreras et al., 2012), ultimately leading to a significant activity of the BK channel. Hence, in cells like neurons, an appreciable open probability of BK channels at physiologically relevant voltages necessarily involves the activation of Ca^2+^ sensors on the gating ring. The allosteric interplays established between the functional and structural modules (VSD-PD, CTD-PD, and CTD-VSD) are key in enabling BK channels to operate over a dynamic wide-range of internal Ca^2+^ and voltage conditions by fine-tuning the channel’s gating machinery. Therefore, understanding the structure-functional bases that underlie the Ca^2+^ and voltage activation mechanisms interrelationship becomes essential.

The voltage dependence of Ca^2+^-dependent gating ring rearrangements (Miranda et al., 2013, 2018) and RCK1 site occupancy (Sweet and Cox, 2008; Savalli et al., 2012; Miranda et al., 2018) as well as the perturbation of VSD movements by Ca^2+^ binding (Savalli et al., 2012) support the idea that the energetic interaction between both specialized sensors may be crucial to favor BK channel activation. The physical CTD-VSD interface has been suggested to provide the structure capable of mediating the crosstalk between these sensory modules and their synergy in activating the pore domain (Yang et al., 2007; Sun et al., 2013; Tao et al., 2017; Zhang et al., 2017). However, the strength of the interaction between voltage and Ca^2+^ sensors and their relevance to BK channel activation is still an unresolved matter (Horrigan and Aldrich, 2002; Carrasquel-Ursulaez et al., 2015). Also, the functional role that plays each of the high-affinity Ca^2+^-binding sites on the CTD-VSD allosteric interaction is an open question. The RCK1 and RCK2 Ca^2+^-binding sites have distinct functional properties conferred by their different molecular structures and relative positions within the gating ring (Wu et al., 2010; Yuan et al., 2010, 2012; Hite et al., 2017; Tao et al., 2017). Thus, the RCK sites differ in their Ca^2+^ binding affinities (Bao et al., 2002; Xia et al., 2002; Sweet and Cox, 2008), divalent cations selectivity (Oberhauser et al., 1988; Schreiber and Salkoff, 1997; Zeng et al., 2005; Zhou et al., 2012), voltage dependence (Sweet and Cox, 2008; Savalli et al., 2012; Miranda et al., 2018) and in their contribution to allosteric gating mechanism (Yang et al., 2010, 2015). In particular, only the RCK1 site appears to be involved in communicating the Ca^2+^-dependent conformational changes towards the membrane-spanning VSD (Savalli et al., 2012; Miranda et al., 2018). Recently, the *Aplysia* BK structure shows that the N-lobe of RCK1 domain is in a non-covalent contact with the VSD and the S4-S5 linker being this RCK1-VSD interaction surface rearranged when comparing the liganded and Ca^2+^-free structures (Hite et al., 2017; Tao et al., 2017). Actually, it has been hypothesized that any Ca^2+^-induced rearrangements of the gating ring should be ultimately transmitted to the pore domain via the VSD (Hite et al., 2017; Zhou et al., 2017). Thus, defining what extent Ca^2+^ binding influences to VSD is crucial in determining how important is the crosstalk between sensors in decreasing the free energy necessary to open the BK channel.

Here, we examined the Ca^2+^-dependence of the VSD activation estimating the allosteric coupling between Ca^2+^ and voltage sensors. By analyzing gating currents under unliganded and Ca^2+^-saturated conditions, we found a strong energetic influence of the Ca^2+^-binding on the voltage sensors equilibrium in an independent manner of the channel opening. These findings point out that a major component in the synergistic Ca^2+^ and voltage activation of BK channels can reside on the sensory domains communication. We also found that the Ca^2+^-dependent behavior of the voltage sensor activation is consistent with an CTD-VSD allosteric coupling that occurs through a concerted interaction scheme where each Ca^2+^-bound to high-affinity sites affect equally all voltage sensors in the BK tetramer. Notably, we found that the two distinct RCK1 and RCK2 Ca^2+^ sensors exert equivalent contributions on VSD via independent allosteric pathways.

## Results

### Allosteric coupling between Ca^2+^-binding and voltage sensor activation is strong

We characterized the effects of Ca^2+^-binding on voltage sensor activation in BK channels by analyzing gating current measured on inside-out patches of *Xenopus laevis* oocyte membrane. Families of gating currents (*I*_*G*_) were evoked at different intracellular Ca^2+^ concentrations ([Ca^2+^]_i_) ranging from 0.1 to 100μM in K^+^-free solution (***Figure 1A***). For all experiments, first we measured *I*_*G*_ in the nominal absence of Ca^2+^ (“zero Ca^2+^” condition), and then we perfused the internal side with solutions containing Ca^2+^ at increasing concentrations. The amount of gating charge displaced (*Q*_C_) at each Ca^2+^ concentration was obtained by fitting the initial part of the ON-gating current decay to a single exponential (fast ON-gating; see *Methods*) and integrating it. In this manner, we determine only the gating charge displaced before the BK channel opening.

**Figure 1.** Effects of Ca^2+^-binding on VSD activation in BK channels. (**A**) Representative gating current (*I*_G_) recordings at different internal Ca^2+^ concentrations (from 0 to 100 μM). *I*_*G*_ were evoked by the indicated voltage protocol of 1 ms duration. (**B**) Gating charge-voltage relationships (*Q*_C_ (*V*)) were obtained by integrating the fast component for each ON *I*_*G*_ trace. Normalized gating charge data (*Q*_C_ (*V*)/*Q*_C, MAX_) (mean ± SEM) were fitted using a single Boltzmann function (solid lines). (**C**) *V*_H_ obtained from the *Q*_C_(*V*) curves as a function of Ca^2+^ concentration (mean ± SEM). At zero” Ca^2+^ condition *V*_H (0 Ca^2+^)_ = 174.5 ± 2.4 mV (*n* = 25), whereas Ca^2+^ binding produce a leftward shift in *V*_H_ (Δ*V*_H_): 0.1 μM Ca^2+^ (Δ*V*_H_ = - 12.1 ± 3.5 mV, *n* = 5); 0.5 μM Ca^2+^ (Δ*V*_H_ = −22.9 ± 1.8 mV, *n* = 5); 1 μM Ca^2+^ (Δ*V*_H_ = −37.1 ± 3.5 mV, *n* = 5); 5 μM Ca^2+^ (Δ*V*_H_ = −58.8 ± 6.7 mV, *n* = 6); 10 μM Ca^2+^ (Δ*V*_H_ = −127.9 ± 13.9 mV, *n* = 6); 100 μM Ca^2+^ (Δ*V*_H_ = −142.6 ± 4.5 mV, *n* = 7). The one-way ANOVA followed by Dunnett’s post-hoc test analysis was used to assess statistical significance of the Ca^2+^-induced shifts in *V*_H_ (\*\*\**p*<0.001).

The increase in internal Ca^2+^ promotes a leftward shift of the *Q*_C_ versus voltage (*Q*_C_(*V*)) curves (***Figure 1B,C***) which indicate that Ca^2+^-binding facilitates the activation of the voltage sensor being more prominent as binding sites occupancy increases. Revealing a strong energetic interaction between both sensors, a significant Ca^2+^-induced shift of voltage sensor activation occurs (Δ*V*_H_ = −142.6 ± 4.5 mV) under Ca^2+^-saturated conditions for high-affinity binding sites (100 μM). Such large shift means that Ca^2+^-binding to the RCK Ca^2+^-binding sites alters the VSD equilibrium promoting a decrease in the free energy 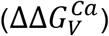 that defines the voltage sensor resting-active (R-A) equilibrium by ~8 kJ/mol 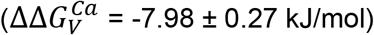.

A visual inspection of the current records indicates that the kinetics of the ON-gating current is not much affected by the concentration of Ca^2+^ present in the internal solution. However, it is apparent that the OFF-gating current is dramatically modified becoming smaller in amplitude and with slower kinetics as the internal Ca^2+^ concentration is increased (***Figure 1A*** and ***Figure 1—figure supplement 1A***). At least two components can be resolved in the OFF gating current decay (***Figure 1—figure supplement 1B,C***), and the relative contribution of the slower component increases as internal Ca^2+^ augmented reflecting an increase of the open probability of channel ***Figure 1—figure supplement 1D,E***). This kinetic behavior recapitulates the effect describe on gating charge displacement as a function of the depolarizing pulse duration (Horrigan and Aldrich, 2002; Contreras et al., 2012; Carrasquel-Ursulaez et al., 2015), and confirm that this phenomenon is associated with the time course of channel opening revealing the allosteric interaction between voltage sensors and the pore gate (Horrigan and Aldrich, 2002).

### Ca^2+^ binding to single α-subunit affects the R-A voltage sensor equilibrium of all four subunits equally

Taking advantage of the dose-dependent effect of Ca^2+^ on voltage sensor activation we investigated the underlying mechanism of the Ca^2+^-voltage sensors communication in the context of the well-established Horrigan-Aldrich (HA) allosteric gating model(Horrigan and Aldrich, 2002). Two different mechanisms were proposed by Horrigan and Aldrich for the interaction between the Ca^2+^-binding sites and voltage sensors. The first mechanism supposes that Ca^2+^ binding to one α-subunit affects the VSD in the same subunit only (Scheme I) (***Figure 2A***), while the second mechanism assumes that the Ca^2+^ binding affects the four VSD equally (Scheme II) (***Figure 2B***). It should be noted that the standard HA model makes two simplifying assumptions by considering a single Ca^2+^-binding site per α-subunit and the Scheme I as the Ca^2+^ binding-VSD interaction mechanism underlying BK channel gating (Horrigan and Aldrich, 2002).

**Figure 2.**
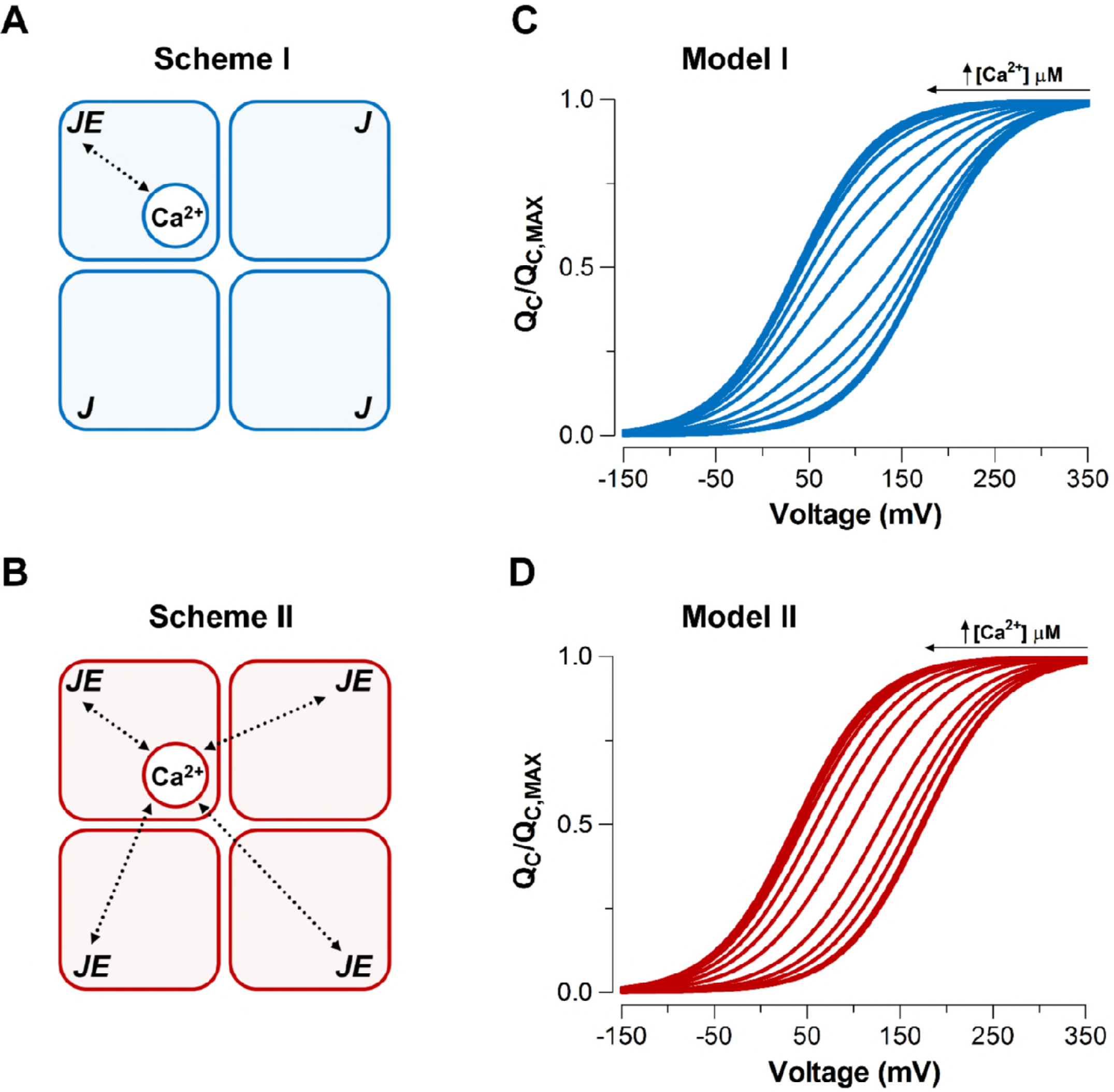
Model-dependent behavior of the *Q*_C_(*V*) curves based on the CTD-VSD interaction mechanisms according to the fractional occupancy of Ca^2+^-binding sites. (**A-B**) Cartoons representing two interaction schemes between voltage sensors and Ca^2+^-binding sites (modify from Horrigan and Aldrich (2002)). The Scheme I (**A**) assume that Ca^2+^-binding only affects the voltage sensor of one α-subunit (*E*_*M*1_). The Scheme II (**B**) predicts that binding of Ca^2+^ to one α-subunit affects VSD in all subunits equally, increasing the voltage sensor equilibrium constants (*J*) *E*_*M*2_-fold in all four subunits (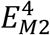, when the four Ca^2+^ sites are occupied). In both schemes, a single Ca^2+^-binding site is considered in each α-subunit. (**C-D**) Predictions of *Q*_C_(*V*) relationships at different internal calcium concentration (from 0 to 10 mM) by two distinctive interaction mechanisms between Ca^2+^-binding sites and voltage sensors (Scheme I and Scheme II), respectively. *Q*_C_(*V*) curves were generated using Equation 4 (blue: Model I) or Equation 6 (red: Model II), and the following set of parameters: *z*_*J*_ = 0.58, *J*_0_ = 0.018, *K*_*D*_ = 11 μM and 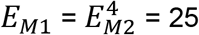.

For a better comprehension, we simulated the normalized *Q*_C_(*V*) curves over a wide range of Ca^2+^ concentrations (from 0 to 10 mM) for each Ca^2+^-VSD interaction scheme (***Figure 2C,D***). Here we assume that the measurement of the fast gating currents captures the charge displaced by R-A transitions and exclude the charge associated with the transition between the activated estates. This assumption is reasonable since the Ca^2+^-binding rate constant estimated for BK is about 10^8^ M^−1^s^−1^ (Hou et al., 2016) implying that at 10 μM internal Ca^2+^ (the highest non-saturating Ca^2+^ concentration tested) the time constant of the Ca^2+^ binding is 1 ms, while the VSD activates with a time constant of ~30 μs at the higher voltage tested (see the *Supplementary Information* for details of the simulations). At extreme conditions of low (0.03 to 0.1 μM) and high (≥100 μM) internal Ca^2+^, the VSD activation behaves in a mechanism-independent manner since all voltage sensors are in the same functional state (unliganded or saturated). However, the distinctive effects on *Q*_C_(*V*) curves at intermediate Ca^2+^ concentrations (1−10 μM) provide useful signatures to distinguish between these two mechanisms. Scheme I predicts two functional states of the VSD depending on occupancy status of the Ca^2+^ site (Ca^2+^ bound and unbound) such that the *Q*_C_(*V*) curve behavior is described by the fractional distribution of the unliganded and Ca^2+^-saturated functional states like an all-or-none allosteric effect (***Figure 2C***; ***Figure 2—figure supplement 1B*** and Equation 4 in *Supplementary Information*). On the contrary, the Ca^2+^-binding effect on the VSD activation according to the Scheme II is characterized by a five-component Boltzmann function (***Figure 2—figure supplement 1C*** and Equation 6 in *Supplementary Information*). Each component represents a single functional state determined by the number of Ca^2+^ bound to the channel (from 0 to 4). In such case, the *Q*_C_(*V*) curves resulting from a distribution of functional states behaves as an equivalent single Boltzmann leftward shifted by an incremental allosteric effect (from *E* to *E*^4^) as the number of Ca^2+^ bound to the channel increases (***Figure 2D***).

To elucidate the mechanism by which Ca^2+^ and voltage sensors interact, we performed fits of the *Q*_C_(*V*) data using the two different models represented in the Scheme I and Scheme II (***Figure 3A,B***). The allosteric factor *E* that accounts for the coupling between the Ca^2+^-binding sites and the voltage sensors was constrained to values calculated from the experimental data of the *Q*_C_(*V*) shift at the Ca^2+^ saturating conditions (100 μM) in relation to the same curve in the absence of Ca^2+^. The *z*_*J*_, *J*_0_ and *K*_*D*_ parameters obtained during the fitting procedure of each model are very similar (***Figure 3C***). The fitted values of the affinity constant (*K*_*D*_ = 3 - 5 μM) agree with previous reports (Cox et al., 1997; Horrigan and Aldrich, 2002) although slightly smaller than those estimated on the closed conformation of the channel (*K*_*D*_ = 11 μM). However, we found that the fit with the Scheme II to the *Q*_C_ (*V*) curves (***Figure 3B***) is better than the fit to the data using Scheme I (***Figure 3A***) as indicated by the Akaike model selection criteria (AIC) (Akaike, 1974). Moreover, Model II generates a *V*_H_-log[Ca^2+^] curve (solid line) that accounts for the dose-response *V*_H_-log[Ca^2+^] experimental data reasonable well (***Figure 3D***). Also, the behavior of *Q*_C_(*V*) curves at intermediate Ca^2+^ concentrations (1−10 μM) is qualitatively consistent with the phenotype exhibit by the Ca^2+^-VSD scheme II (***Figure 2D*** and ***Figure 3B***). Thus, the experimental dose-dependent effect of Ca^2+^ on voltage sensor activation reveals that Ca^2+^-binding to a single α-subunit of BK channels increases *E*-fold the equilibrium constant *J* that defines the equilibrium between resting and active conformations of the voltage sensors in all four subunits.

**Figure 3.** Dose-dependent effect of Ca^2+^ on voltage sensor activation is predicted by a Ca^2+^-VSD interaction mechanism defining that Ca^2+^-binding equally affects the VSD in all four α-subunits. (**A-B**) The experimental *Q*_C_(*V*) data were fitted with the two possible allosteric interaction mechanisms between voltage and calcium sensors described by the Scheme I and Scheme II. The blue and red lines represent the global fits by Model I and Model II, respectively. The allosteric factor *E* (*E*_*M*1_ and *E*_*M*2_) was constrained to the value obtained from the individual fitting of the *Q*_C_(*V*)/*Q*_C, MAX_ curves at 0 and 100 μM Ca^2+^ (experimental *E* (*E*_*exp*_) equal to 26.4). The *z*_*J*_, *J*_0_ and *K*_*D*_ parameters were allowed to vary freely. (**C**) Parameters for the best fits of the *Q*_C_(*V*) data. Note that the allosteric factor *E* for Model I (*E*_*M*1_) and Model II (*E*_*M*2_) have different interpretations, being *E*_*M*2_ = *E*_*exp*_ whereas 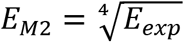 given that the four voltage sensor will be altered in 2.3-fold (*E_M2_* = 2.27) with each additional Ca^2+^ bound. Based on the Akaike Information Criterion (AIC), the best model fit to the *Q*_C_(*V*, [Ca^2+^]) data is achieved using the Ca^2+^-VSD interaction scheme described by Model II. (**D**) The Ca^2+^-dependence of *V*_H_-*Q*_C_(*V*) curves are superimposed with the predicted *V*_H_ by Model II (red line).

### High-affinity Ca^2+^-binding sites in RCK1 and RCK2 domains contribute equally to the allosteric coupling between Ca^2+^ and voltage sensors

Under physiological conditions, the RCK1 and RCK2 high-affinity Ca^2+^-binding sites are responsible by all calcium sensitivity of the activation of BK channel (Schreiber and Salkoff, 1997; Bao et al., 2002, 2004; Xia et al., 2002). However, distinct physiological roles of the RCK1 Ca^2+^-sensor and Ca^2+^ bowl may be based in their functionally and structurally distinctive properties (Zeng et al., 2005; Sweet and Cox, 2008; Yang et al., 2010; Savalli et al., 2012; Tao et al., 2017). Below, we asked what the energetic contribution to VSD equilibrium is of the two high-affinity Ca^2+^-binding sites contained in the RCK1 and RCK2 domains.

To elucidate the effect of each Ca^2+^-sensor on the VSD activation we used mutations that selectively and separately abolish the function of the two differents RCK Ca^2+^-sites. Disruption of the RCK1 Ca^2+^-sensor by the double mutant D362A/D367A (Xia et al., 2002) reduces significantly (48%, Δ*V*_H (D362A/D367A)_ = −74.9 ± 4.7 mV) the leftward shifted of the *Q*_C_(*V*) curves at 100 μM Ca^2+^ compared with the wild-type (WT) BK channel (***Figure 4A,C***). We also examined the effect of the mutant M513I (Bao et al., 2002) which have been shown to eliminate the Ca^2+^ sensitivity derived from the RCK1 site (Bao et al., 2002, 2004; Zhang et al., 2010). In this mutant, the 100 μM Ca^2+^-induced shift in *V*_H_ of VSD activation curve is also considerably smaller relative to WT (about 54%, Δ*V*_H (M513I)_ = - 65.4 ± 2.6 mV) (***Figure 5***). Therefore, both mutations affect the Ca^2+^-induced enhancement of activation of voltage sensor very similarly through the RCK1 site (***Figure 5C***), although their mechanisms action could be quite different. The M513 residue appears to participate in the stabilization of the proper conformation RCK1 Ca^2+^-site whereas D367 is a key residue in the coordination of Ca^2+^ ion (Wu et al., 2010; Zhang et al., 2010; Tao et al., 2017). On the other hand, neutralization of the residues forming part of the Ca^2+^ bowl (Schreiber and Salkoff, 1997) (5D5A mutant, see *Methods*) on the RCK2 domain decreases the leftward shift of the *Q*_C_(*V*) curve when Ca^2+^ is increased to 100 μM by approximately 54% (Δ*V*_H (5D5A)_ = −65.7 ± 4.7 mV) (***Figure 4B, D***). Surprisingly, the effect of Ca^2+^ binding on Δ*V*_H_ from each high-affinity Ca^2+^ site is roughly half relative to WT channels with both intact sites (***Figure 4E***). Therefore, both high-affinity Ca^2+^-binding sites contribute equally to decrease the free energy necessary to activate the VSD. Thus, the change of free energy of the resting-active equilibrium of the voltage sensor in response to Ca^2+^-binding at RCK2 site is ~ −4 kJ/mol 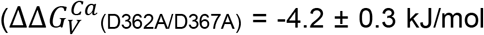 and 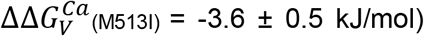 (***Figure 4C*** and ***Figure 5C***). In the same way, the occupation of the RCK1 Ca^2+^-binding site decreases the free energy necessary to activate the VSD in −3.8 ± 0.4 kJ/mol 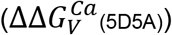. Remarkably, these findings reveal an additive effect of Ca^2+^-binding to the RCK1 and Ca^2+^ bowl sites on the VSD activation which suggest independent allosteric pathways through which they exert their modulation on the VSD.

**Figure 4.** The high-affinity Ca^2+^-binding sites contribute equally to allosteric coupling between calcium and voltage sensors in BK channels. (**A-B**) Representative gatingcurrent (*I*_*G*_) recordings at 0 and 100 μM of [Ca^2+^]_i_ for the RCK1 site mutant (D362A/D367A) and the RCK2 site mutant (5D5A), respectively. (**C-D**) Gating charge-voltage curves (*Q_c_(V)*) were obtained at 0 Ca^2+^ (open symbols) and 100 μM Ca^2+^ (filled symbols) for D362A/D367A and 5D5A mutants, respectively. Boltzmann fitting to the experimental data (mean ± SEM) is indicated by solid lines (μ_H_(D362A/D367A) = 178.0 ± 2.7 mV, *n* = 12 and *V*_H_(5D5A) = 176.4 ± 4.6 mV, *n* = 17 at “zero” Ca^2+^; *V*_H_(D362A/D367A) = 104.2 ± 7.3 mV, *n* = 7 and V_H_(5D5A) = 110.8 ± 6.7 mV, *n* = 6 at 100 μM Ca^2+^). For comparison, all *Q*_C_(*V*) plots include the Boltzmann fit of the *Q*_C_(*V*) curves for WT at 0 Ca^2+^ (dashed black line) and 100 μM Ca^2+^ (solid black line). (**e**) Quantification of the *V_H_* shift (Δ*V*_H_) in the *Q*_C_(*V*) curves and the free energy change 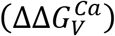 induced by 100 μM Ca^2+^. The non-parametric *t*-test was used to evaluate statistical significances between WT BK channel and the RCK sites mutants (\*\*\**p*<0.001; ns: non-significant).

**Figure 5.**
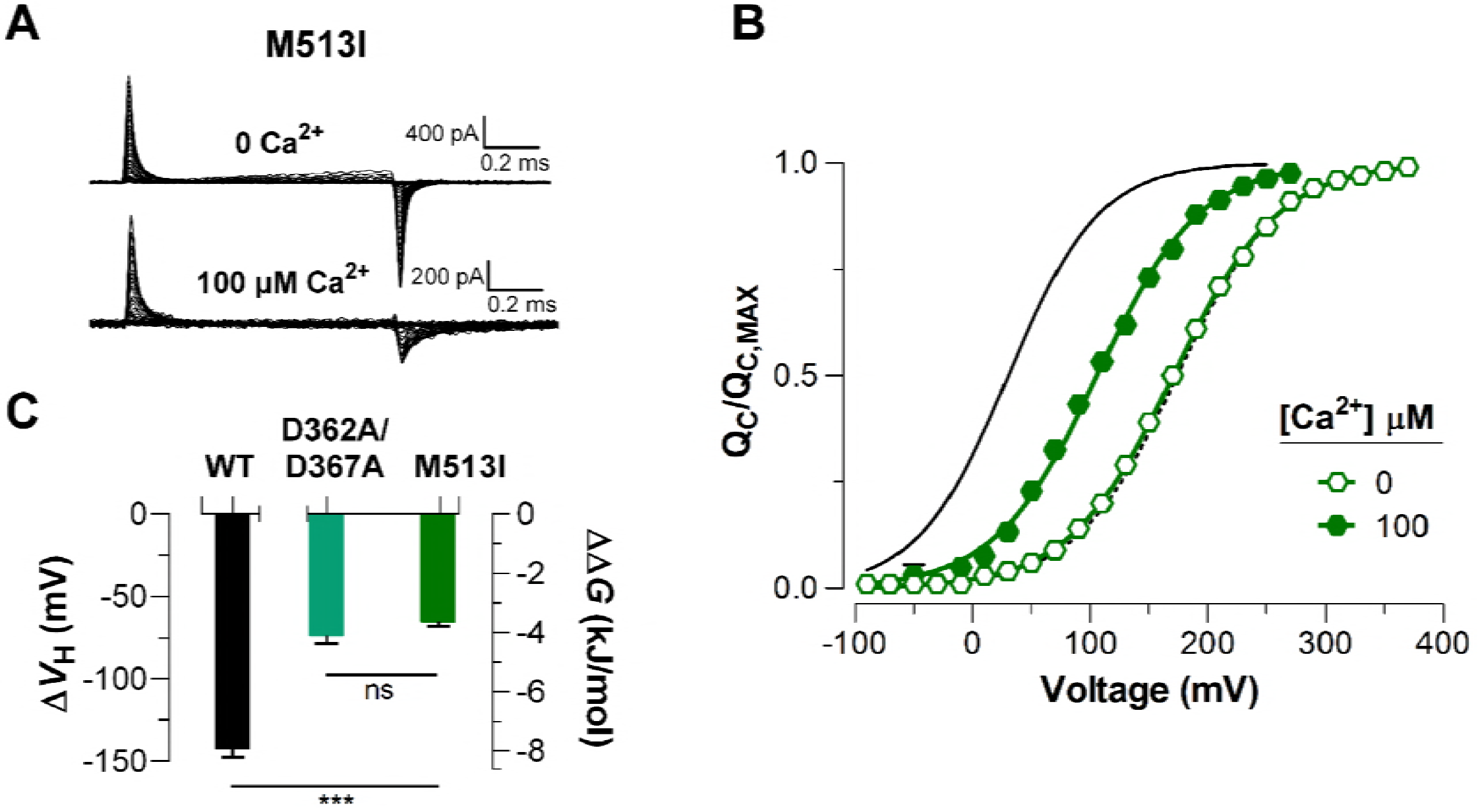
Mutations abolishing Ca^2+^-sensing by the RCK1 binding-site reduce the Ca^2+^-induced effect on voltage sensors activation similarly. (**A**) Representative gating current (I_G_) recordings at 0 and 100 μM of [Ca^2+^]_i_ for the RCK1 site mutant M513I. (**B**) Gating charge-voltage curves *Q*_C_(*V*) were obtained at 0 Ca^2+^ and 100 μM Ca^2+^ (open and filled symbols) for the M513I mutant. Boltzmann fitting to the experimental data (mean ± SEM) is indicated by solid lines (*V*_H_(M513I) = 170.4 ± 4.4 mV, *n* = 17 at “zero” Ca^2+^ and *V*_H (M513I)_ = 105.0 ± 6.3 mV, *n* = 4 at 100 μM Ca^2+^). For comparison, the *Q*_C_(*V*) plot includes the Boltzmann fit of the *Q*_C_(*V*) curves for WT at 0 Ca^2+^ and 100 μM Ca^2+^ (dashed and solid black line). (**C**) Quantification of the *V_H_* (Δ*V*_H_) shift in the *Q*_C_(*V*) curves and the free energy change induced by 100 μM Ca^2+^ 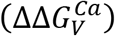. The non-parametric t-test was used to compare statistical significances between WT BK channel and the RCK1-site mutants (***p<0.001; ns: non-significant).

Taking these results into account, we expanded the Ca^2+^-VSD interaction model described by Scheme II considering the energetic contribution of the two kinds of Ca^2+^ sensors on the VSD per α-subunit (*E*_*WT*_ = *E*_*S*1_ * *E*_*S*2_) (***Figure 2—figure supplement 1E***). As described in the above model fittings, the allosteric factors *E* of each one RCK1 and RCK2 sites (*E*_*S*1_ and *E*_*S*2_) were constrained to values equivalent to the Ca^2+^-induced energetic perturbations of the voltage sensor equilibrium for the 5D5A and D362A/D367A mutants, respectively. However, the inclussion of the two Ca^2+^ sensors in the Ca^2+^-VSD interaction model does not produce better fits to *Q*_C_(*V*, [Ca^2+^]) according to the AIC criteria (***Table 1*** and ***Figure 3B***), the estimated *K*_*D*_ parameters for each Ca^2+^-binding sites (*K*_*D*1_ = 15.6 μM and *K*_*D*2_ = 1.9 μM) by the experimental data fitting agrees very well with the apparent Ca^2+^ affinities previously reported in the literature (Bao et al., 2002; Xia et al., 2002; Sweet and Cox, 2008). Interestingly, modest positive cooperativity (*G* = 2.6) between the two Ca^2+^-binding sites located in the same α-subunit is required to achieve a good estimation of the *K*_*D*_ parameters (***Table 1***), where the Ca^2+^ bowl site has an affinity for Ca^2+^ about 8fold greater than does the RCK1 Ca^2+^-sensor.

**Table 1.**
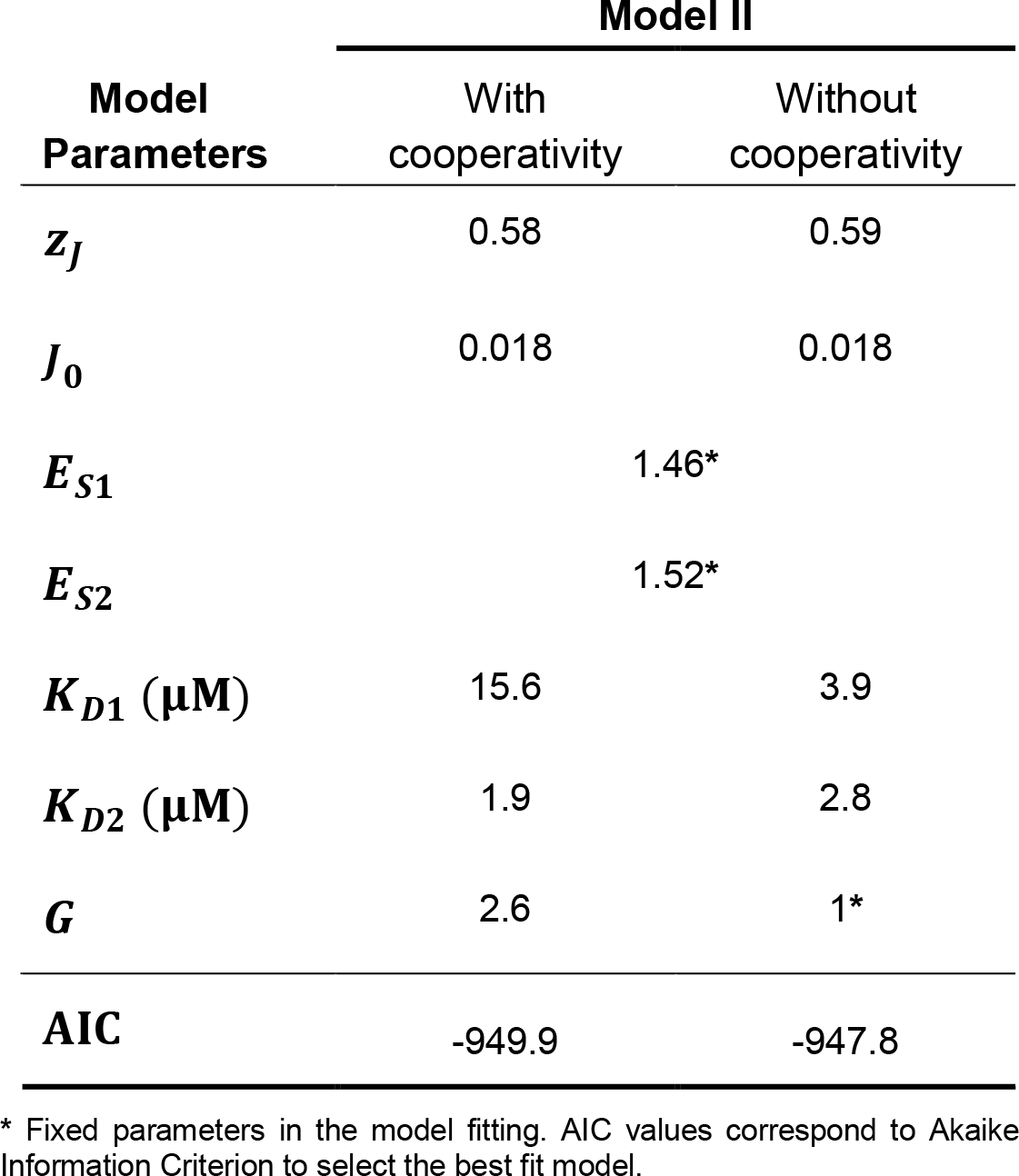
Parameters for the extended Ca^2+^-VSD interaction model (Model Il) including two high-affinity Ca^2+^-binding sites per α-subunit.

## Discussion

Recent insights into a major interplay between voltage- and Ca^2+^-sensing modules in the BK channel are supported by functional and structural studies (Yuan et al., 2010; Savalli et al., 2012; Miranda et al., 2013, 2016, 2018; Carrasquel-Ursulaez et al., 2015; Hite et al., 2017; Tao et al., 2017; Zhang et al., 2017), offering a new perspective in our understanding of its multimodal gating mechanism. However, the CTD-VSD allosteric coupling as well its molecular nature has yet to be firmly established since their direct assessment is subject to great experimental challenges. Based on the functional independence of the distinct structural domains (PD, CTD, and VSD), the energetic relationship between the sensory modules can be directly defined comparing the voltage sensor equilibrium change at extreme Ca^2+^ stimulus conditions limiting the status of the Ca^2+^-binding sites to two well-defined configurations: unliganded and saturated (Horrigan and Aldrich, 2002).

Using this approach, this work straightforwardly establishes that Ca^2+^-binding to high-affinity sites make a significant and direct energetic contribution to the equilibrium of the resting-activated transition (R-A) of the VSD facilitating their activation (Δ*V*_H_ = −142.6 ± 4.5 mV and 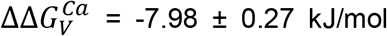). This result resolves a previous debate regarding to the magnitude of the Ca^2+^-driven shift of the *Q*_C_(*V*) curve, because it has been reported a similar leftward shift (Δ*V*_H_ = −140 mV and 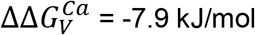; at [Ca^2+^]_i_ = 100 μM) (Carrasquel-Ursulaez et al., 2015) and a smaller leftward shift (Δ*V*_H_ = −33 mV and 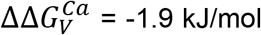; at [Ca^2+^]_i_ = 70 μM) (Horrigan and Aldrich, 2002) at saturating Ca^2+^ concentration. The reason for the contradictory findings is not clear to us; since we used a similar experimental approach. Even if we assumed that the calcium effect on VSD is underestimated at 70 μM Ca^2+^ (Horrigan and Aldrich, 2002) compared to 100 μM as saturating condition of the binding sites, we observed a significative greater effect in Ca^2+^ concentrations (1, 5 and 10 μM, ***Figure 1C***) where less than 50% of the Ca^2+^ sensors are occupied (*K*_*D*_ = 11 μM (Cox et al., 1997; Horrigan and Aldrich, 2002)).

Fluorescence studies that optically track the motions of the voltage sensor or gating ring provide two lines of evidence that support these findings. First, conformational rearrangements of the voltage sensors detected using voltage-clamp fluorometry can be provoked by Ca^2+^-binding to the high-affinity sites. The sudden rise of intracellular [Ca^2+^] by UV flash induced-photolysis of caged Ca^2+^ prompts a leftward shift in both conductance-voltage (*G*(*V*)) and fluorescence-voltage (*F*(*V*)) relationships. These results suggest that functional activation of the gating ring is propagated to VSD leading to structural perturbations of voltages sensors, thereby favoring its active conformation (Savalli et al., 2012). Second, the structural rearrangement of gating ring in response to Ca^2+^ has a voltage dependence (Miranda et al., 2013, 2018) attributable to the voltage sensor operation. The origin of these voltage-dependent motions has been recently established via modifications on the voltage-sensing function of the BK channel using the patch-clamp fluorometry technique (Miranda et al., 2018). Both the charged residue mutations on the S4 transmembrane segment (R210, R213, and E219) and the coexpression of β1-subunit with BKα channel modify the conformational changes of the gating ring triggered by depolarization in correspondence to the observed *G*(*V*) shift for these channel constructs. In contrast, perturbations of pore opening equilibrium like the F315A mutation or the assembly of BKα channel with γ1-subunit does not modify on the voltage-dependent reorganization of the gating ring (Miranda et al., 2018).

Mechanistically and in a channel opening-independent fashion, how does the CTD-VSD coupling occur? Taking into account the homotetrameric configuration of the BK channel, Horrigan and Aldrich (Horrigan and Aldrich, 2002) defined the general gating scheme of BK channel considering the simplest CTD-VSD interaction model in which voltage sensors and Ca^2+^-binding sites solely interact within the same subunit. However, the VSD movement at non-saturating Ca^2+^ conditions which entail distinct functional states of the Ca^2+^ sites (unliganded and liganded), unveiled that the standard HA model can not explain the mechanistic interaction governing the allosteric coupling between the Ca^2+^ and voltage sensors. Given that Ca^2+^-binding will influence only a fraction of voltage sensors, Scheme I would evidence *Q*_C_(*V*) curves characteristic of an all-or-none model showing two well-distinguishable components Boltzmann that correspond to Ca^2+^-affected and unaffected VSD fractions (***Figure 3A***). Conversely, an energetic effect of each Ca^2+^-site on all the voltage sensors of the tetramer would lead to an equivalent functional status of each VSD, so that the *Q*_C_(*V*) curves behaving in an incremental shifted fashion as increasing the fractional occupancy of the Ca^2+^ sites. The VSD and Ca^2+^ sites interacting in such a fashion (Scheme II) reproduce reasonably well the behavior of the Ca^2+^-dependent gating charge movement observed in our experiments (***Figure 3B***). This concerted CTD-VSD communication may underlie a mechanism analogous to the mechanical strategy of interaction between the homooctameric ring of RCK domains and the pore module described for bacterial K^+^ channels (Jiang et al., 2002; Ye et al., 2006; Lingle, 2007; Pau et al., 2011; Smith et al., 2012, 2013). Both in MthK and BK channels, the Ca^2+^-site occupancy triggers a conformational change conveying to a symmetric overall rearrangement of the cytosolic tetrameric structure that finally is propagated to the transmembrane regions (TMD) via C-linker and in the BK channel also via the protein-protein interfaces between the gating ring and the TMD (Jiang et al., 2002, 2003; Ye et al., 2006; Yuan et al., 2010, 2012; Pau et al., 2011; Smith et al., 2012; Tao et al., 2017). Consequently, we can speculate that each Ca^2+^-binding event produces a gradual conformational expansion of the gating ring affecting the four voltage sensors in each step through the progressive perturbations within the protein-protein interfaces.

As mentioned above, the communication pathway through which the Ca^2+^-driven conformational changes are propagated to the voltage sensors appears to critically reside on the CTD-VSD interface that involves non-covalent interactions between RCK1 N-lobe and S0-S4 transmembrane segments (Yang et al., 2007, 2008, 2010; Sun et al., 2013; Hite et al., 2017; Tao et al., 2017). Scanning mutagenesis of RCK1-N terminal subdomain indicated that residues on the βA-αC region are involved into the allosteric connection of the Ca^2+^-dependent activation mediated by RCK1 site occupancy but not to the Ca^2+^ bowl (Yang et al., 2010). In line with this study, the selective activation of the RCK1 domain was identified to be responsible for the Ca^2+^-induced VSD rearrangement (Savalli et al., 2012) and the voltage dependence of the Ca^2+^-driven motions of gating ring (Miranda et al., 2016, 2018), suggesting that CTD-VSD allosteric coupling is primarily determined by the RCK1 site. However, our results are inconsistent with this picture. The constructs D362A/D367A and 5D5A (D894A-D898A) selectively impaired the Ca^2+^-sensitivity of the RCK1- and RCK2-sensors, respectively, by neutralization of residues that are involved in contributing to Ca^2+^-coordination (Zhang et al., 2010; Tao et al., 2017). Comparing the fast gating charge movement at 0 Ca^2+^ and saturating Ca^2+^ conditions reflects that the energetic effect of Ca^2+^-binding on voltage sensor equilibrium is practically identical (~ −4 kJ/mol) for either the D362A/D367A mutations or 5D5A mutations (***Figure 4***). Thus, our findings establish that the RCK2-driven contribution to CTD-VSD energetic coupling is quite similar to the RCK1-driven contribution. The functional role of the RCK2-sensor on Ca^2+^-sensitivity of VSD activation was further corroborated using the M513I mutation (***Figure 5***). This point mutation hinders the Ca^2+^-dependent activation associated with the RCK1-sensor presumably by disrupting the structural integrity of the binding site and the transduction via through the βA-αC region (Zhang et al., 2010). Thus, another residue involved in the BK Ca^2+^-dependent activation mediated by the RCK1 Ca^2+^-binding site but not forming part of the site itself decreases the *Q*_C_(*V*) leftward shift almost in the same amount as it does the D362A/D367A mutant.

Interestingly, beyond the energetic contribution of each RCK site to the voltage sensor equilibrium is the same, its addition mimics the VSD Ca^2+^-sensitivity of the fully occupied sites. These findings remind us of early reports showing that each RCK site mutant shifts the Ca^2+^-dependent *G*(*V*) by approximately half relative to WT channels (Bao et al., 2002; Xia et al., 2002). Our results suggest an autonomy of the two RCK-sensors indicating independent allosteric pathways through which they exert their modulation on the VSD but does not discard some cooperativity effect between them. Indeed, various lines of evidence indicate albeit modest a cooperativity between the two high-affinity Ca^2+^-binding sites although their nature is still unclear (Qian et al., 2006; Sweet and Cox, 2008; Savalli et al., 2012). Intra and intersubunit structural connectivity support the putative cooperative interactions between the Ca^2+^ sensors at the gating ring (Yuan et al., 2012; Hite et al., 2017). Actually, a recently functional study of the intrasubunit connections between the RCK1 site and Ca^2+^ bowl (R514-Y904/E902 interactions) has shown that such connections are potential candidates of the structural determinants underlying to a cooperative mechanism between the RCK1- and RCK2-sensor involving either to preserve the integrity of RCK1 Ca^2+^-binding site or the allosteric propagation pathway towards transmembrane domains (Kshatri et al., 2018). On the basis of the cryo-EM structure of *Aplysia californica* BK channel, Hite *et al.* (Hite et al., 2017) proposed that there should be a positive cooperativity of the Ca^2+^-binding at RCK1 site and Ca^2+^ bowl since the Ca^2+^-induced conformational change of the RCK1-N lobes from closed to open configuration depends on functional state (unliganded and liganded) of both RCK sites.

Our analysis based on the CTD-VSD interaction model predicted a small and positive cooperative relation (*G* = 2.6) among the two high-affinity Ca^2+^ sites within the same α-subunit, which has been suggested by an earlier study (Qian et al., 2006). It is noteworthy that *K*_*D*_ parameters achieved for each Ca^2+^-binding sites (*K*_*D*1_ = 15.6 μM and *K*_*D*2_ = 1.9 μM) by the experimental data fitting agrees very well with the apparent Ca^2+^ affinities previously reported in the literature (Bao et al., 2002; Xia et al., 2002; Sweet and Cox, 2008). Together, all this new information recapitulate a more relevant functional role of the cooperative interactions between RCK sensors within the same subunit on Ca^2+^-dependent activation of the channel (Qian et al., 2006).

In conclusion, our results depict a remarkable, and direct energetic direct interplay between the specialized sensory modules (VSD and CTD). Our findings together with the emerging structural-functional information establish a new paradigm about how the stimuli integration (depolarization and intracellular Ca^2+^) modulates the BK channel activation and its relevance within a physiological context. Notable and unexpected is the equivalent role of the distinct ligand-binding sites at the cytosolic domain to the allosteric regulation on voltage sensing. Additional studies to discern the molecular bases underlying in the Ca^2+^ and voltage propagation pathways and the cooperative interactions of the RCK1 and RCK2 regulatory domains may provide new clues about the dual gating mechanism of BK channel.

## Methods

### Channel Expression

*Xenopus laevis* oocytes were used as a heterologous system to express BK channels. The cDNA coding for the human BK α-subunit (U11058) was provided by L. Toro (University of California, Los Angeles, CA). The cDNA coding for independent mutants of each two high-affinity Ca^2+^ site from BK channel, the double mutant D362A/D367A (Xia et al., 2002) and the mutant M513I (Bao et al., 2002) in the RCK1 Ca^2+^-binding site and the mutant 5D5A (Schreiber and Salkoff, 1997) (D894A/D895A/D896A/D897A/D898A) in the RCK2 Ca^2+^-binding site or calcium bowl, were kindly provided by M. Holmgren (National Institutes of Health, Bethesda, MD). The cRNA was prepared by using mMESSAGE mMACHINE (Ambion) for *in vitro* transcription. *Xenopus laevis* oocytes were injected with 50 ng of cRNA and incubated in an ND96 solution (in mM: 96 NaCl, 2 KCl, 1.8 CaCl_2_, 1 MgCl_2_, 5 HEPES, pH 7.4) at 18°C for 4−8 days before electrophysiological recordings.

### Electrophysiological recordings

All recordings were made by using the patch-clamp technique in the inside-out configuration. Data were acquired with an Axopatch 200B (Molecular Devices) amplifier and the Clampex 10 (Molecular Devices) acquisition software. Gating current (*I*_*G*_) records were elicited by 1-ms voltage steps from −90 to 350 mV in increments of 10 mV. Both the voltage command and current output were filtered at 20 kHz with 8-pole Bessel low-pass filter (Frequency Devices). Current signals were sampled with a 16-bit A/D converter (Digidata 1550B; Molecular Devices), using a sampling rate of 500 kHz. Linear membrane capacitance and leak subtraction were performed based on a P/4 protocol (Armstrong and Bezanilla, 1974).

Borosilicate capillary glasses (1B150F−4, World Precision Instruments) were pulled in a horizontal pipette puller (Sutter Instruments). After fire-polished, pipette resistance was 0.5−1 MΩ. The external (pipette) solution contained (in mM): 110 tetraethylammonium (TEA)-MeSO_3_, 10 HEPES, 2 MgCl_2_; pH was adjusted to 7.0. The internal solution (bath) contained (in mM): N-methyl-D-glucamine (NMDG)-MeSO_3_, 10 HEPES, and 5 EGTA for “zero Ca^2+^” solution (~0.8 nM, based on the presence of ~10 μM contaminant [Ca^2+^] (Cui et al., 1997). For test solutions at different Ca^2+^ concentrations (0.1−100 μM), CaCl_2_ was added to reach the desired free [Ca^2+^], and 5 mM EGTA (0.1−0.5 μM) or HEDTA (1−10 μM) was used as calcium buffer. No Ca^2+^ chelator was used in 100 μM free Ca^2+^ solutions. Free calcium concentration was estimated using the WinMaxChelator Software and checked with a Ca^2+^-electrode (Hanna Instruments). All experiments were performed at room temperature (20−22 °C). To measure *I*_*G*_ at different Ca^2+^ concentrations in the same oocyte, the patch was excised and washed with an appropriate internal solution at least 10 times the chamber volume.

### Data Analysis

All data analysis was performed using Clampfit 10 (Molecular Devices), Matlab (MathWorks) and Excel 2007 (Microsoft). The first 50−100 μs of the ON-gating currents were fitted to a single exponential function and the area under the curve was integrated to obtain the charge displaced between closed states (*Q*_C_) (Horrigan and Aldrich, 1999, 2002; Contreras et al., 2012; Carrasquel-Ursulaez et al., 2015). *Q*_C_(*V*) data for each [Ca^2+^]_i_ were fitted using a Boltzmann function: 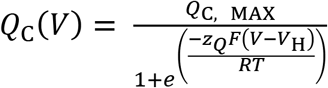 where *Q*_C, MAX_ is the maximum charge, *z*_*Q*_ is the voltage dependency of activation, *V*_H_ is the half-activation voltage, *T* is the absolute temperature (typically 295 K), *F* is the Faraday’s constant, and *R* is the universal gas constant. *Q*_C, MAX_, *V*_H_, and *z*_*Q*_ were determined using least square minimization. *Q*_C_(*V*) curves were aligned by shifting them along the voltage axis by the mean Δ*V* = (〈*V*_H_〉− *V*_H_) to generate a mean curve that did not alter the voltage dependence (Horrigan and Aldrich, 1999). All error estimates are SEM.

The Ca^2+^-induced effect on VSD activation was quantified as the *V*_H_ shift relative to “zero” Ca^2+^ condition: 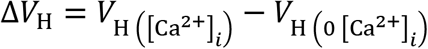. For wild-type (WT) BK channel and the RCK Ca^2+^-sensor mutants (D362A/D367A, M513I and 5D5A), the energetic contribution of Ca^2+^-binding on resting-active (R-A) equilibrium of the voltage sensor was calculated as changes in Gibbs free energy of VSD activation induced by 100 μM Ca^2+^: 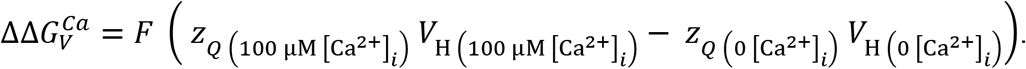.

### Model fitting

We fit the *Q*_C_(*V*,[Ca^2+^]) experimental data using two distinct interaction mechanism between Ca^2+^-binding sites and voltage sensor (see Scheme I and Scheme II in the ***Figure 2A,B***) within the framework of Horrigan-Aldrich (HA) general allosteric model (Horrigan and Aldrich, 2002). Assumptions and considerations for the equations that describe each one of the Ca^2+^-VSD interaction schemes are given in the *Supplementary Information.* In terms of the HA allosteric mechanisms, the voltage sensor R-A equilibrium is defined by the equilibrium constant *J* according to the relation 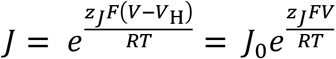, where *J*_0_ is the zero voltage equilibrium constant and *z*_*J*_ the gating charges displacement per voltage sensor. In this fashion, the fraction of the total charge displaced essentially between closed states, (*Q*_C_(*V*)/*Q*_C, MAX_) in the absence of calcium can be written as: 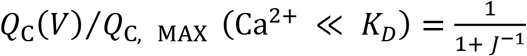, where *K*_*D*_ is the dissociation constant of the high-affinity calcium-binding site with all voltage sensors at rest and the channel closed. In the presence of saturating Ca^2+^ (100 μM), the equilibrium of the R-A transition *J* becomes amplified by the allosteric factor *E*, which defines the coupling between Ca^2+^ binding sites and voltage sensors, being 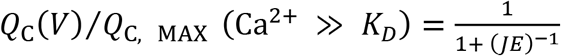 and 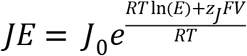. The *Q*_C_(*V*)/*Q*_C, MAX_ measured in the presence of high [Ca^2+^] and “zero Ca^2+^” condition at the same voltage (so that*J* be canceled out) but in the limit where voltage where *J* ^−1^ ≫ 1 is:

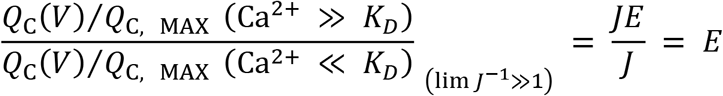

Thus, the Gibbs free energy perturbation of the voltage sensor R-A equilibrium when the high-affinity binding sites are approximately 100% occupied by Ca^2+^ (100 μM) is a straightforward measure of the allosteric factor *E*: 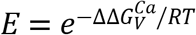

Based on these conditions, the allosteric parameter *E* values were calculated and introduced in each of the two Ca^2+^-VSD interaction models as a fixed parameter. Once *E* was obtained, the families of *Q*_C_(*V*, [Ca^2+^]) curves were simultaneously fitted to the model equations (Equation 3 and Equation 6) (see *Supplementary information*) by minimizing least-squares estimating the *z*_*J*_, *J*_0_ and *K*_*D*_ parameters for each model. To select the better Ca^2+^-VSD interaction scheme that describes the experimental data, the model fits were compared according to their Akaike Information Criterion (AIC) (Akaike, 1974) values, calculated as AIC = 2*p* − 2 ln(*L*), where *p* is the number of free parameters and ln(*L*) is the maximum log-likelihood of the model. The best model fitting is that achieving the lowest AIC values. Minimum AIC values were used as model selection criteria.

The best model fit of the Ca^2+^-VSD interaction scheme was extended including two high-affinity Ca^2+^-binding sites per α-subunit (***Figure 2—figure supplement 2D,E***). The contribution of each Ca^2+^-binding site to the free energy of the voltage sensor equilibrium may be split in two, such as 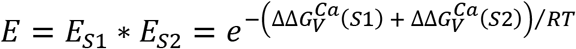, where *E*_*S*1_ and *E*_*S*2_ are the allosteric factor *E* for the RCK1 and RCK2 sites. Thus, for the global fit of the *Q*_C_(*V*, [Ca^2+^]) curves, we constrained the allosteric parameter *E*_*S*1_ and *E*_*S*2_ obtained experimentally for the RCK2 Ca^2+^-sensor mutant (5D5A) and RCK1 Ca^2+^-sensor mutant (D362A/D367A), respectively, as described above. The rest of the parameters *z*_*J*_, *J*_0_, *K*_*D*1_, *K*_*D*2_, and *G*, where *K*_*D*1_ and *K*_*D*2_ are the dissociation constants of the RCK1 and RCK2 sites and *G* is a cooperativity factor between the two sites within the same α-subunit of the BK channel, were allowed to vary freely.

## Acknowledgments

We thank Mrs. Luisa Soto (University of Valparaiso) for excellent technical assistance. This research was supported by FONDECYT Grant No. 1150273 and AFOSR No. FA9550−16−1−0384 to R.L.; CONICYT-PFCHA Doctoral fellowships to Y.L.C.; FONDECYT Grant No. 1180999 to K.C. The Centro Interdisciplinario de Neurociencia de Valparaiso is a Millennium Institute supported by the Millennium Scientific Initiative of the Chilean Ministry of Economy, Development, and Tourism (P029−022-F).

## Supplementary Information

### Assumptions and model predictions

We assume that the four voltage sensors act independently transiting between two states, resting (R) and active (A), governed by the voltage-dependent equilibrium constant *J*. The R-A equilibrium is displaced toward the active state by membrane depolarization generating a fast gating charge movement (*Q*_C_) before channels opening. Additionally, the Ca^2+^-binding to high-affinity sites shifts the voltage sensor equilibrium toward their active configuration through an allosteric coupling described by the factor *E* (***Figure 2—figure supplement 1A***). By assuming the simplified standard model for the BK channels (Horrigan and Aldrich, 2002), where each α-subunit has a single Ca^2+^-binding site, we established the possible states and their connections through which each voltage sensor transit in presence of Ca^2+^ (***Figure 2—figure supplement 1B;C***) following the CTD-VSD interaction mechanisms described by the Scheme I and Scheme II (***Figure 2A,B***).

For Scheme I, in which Ca^2+^-binding sites and voltage sensors can only interact within the same α-subunit, the activation of each VSD can occur through the R_0_-A_0_ or R_1_-A_1_ transitions according to the functional state of the Ca^2+^ site (unbound or Ca^2+^ bound). The equilibrium of such transitions is governed by *J* or *JE*_*M*1_, respectively (***Figure 2—figure supplement 2B***). In the case of Scheme II, in which binding of Ca^2+^ to a single α-subunit affects the four voltage sensors equally, the R-A equilibrium of each VSD would be affected by the number of Ca^2+^ bound in the channel (0−4) depicted in the model (Model II) as five possible R-A transitions. According to this model, the *J* constant increase *E*_*M*2_-fold for each occupied Ca^2+^ site (***Figure 2—figure supplement 1C***). For both schemes, the horizontal transitions R-R and A-A represent the Ca^2+^-binding equilibrium (*K* or *KE*) when the VSD is in the resting or active conformation, respectively. The *K* equilibrium constant is defined as the bound/unbound probability ratio for each Ca^2+^-binding site and depends on Ca^2+^ concentration ([Ca^2+^]) and the Ca^2+^ dissociation constant (*K*_*D*_): *K* = [Ca^2+^]/*K*_*D*_.

Here, we assume that the voltage sensor movement at ON-gating currents is in equilibrium relative to the binding of Ca^2+^. The assumption is reasonable since the Ca^2+^-binding rate constant estimated for BK channel is about 10^8^ M^−1^s^−1^ (Hou et al., 2016) implying that at 10 μM internal Ca^2+^ the association time constant is 1 ms. Thus, Ca^2+^ binding at this Ca^2+^ concentration proceeds at a pace about 33-fold slower than the voltage sensor movement (~30 μs). Based on this consideration, the R-A transitions in the models would be predominant transitions whose proportion will be determined by the [Ca^2+^] and *K*_*D*_. Therefore, predictions of the *Q*_C_(*V*) curves at different Ca^2+^ concentrations for Model I and Model II were based on a given fractional occupancy of Ca^2+^ sites established by the probability of Ca^2+^ bound (*b*) and unbound (1 − *b*) for each Ca^2+^-sensor, and the energetic contribution to VSD equilibrium.

Simulations of the *Q*_C_(*V*) curves using the Scheme I (Model I) were obtained using the equation

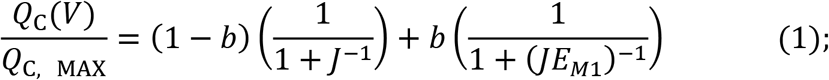

where

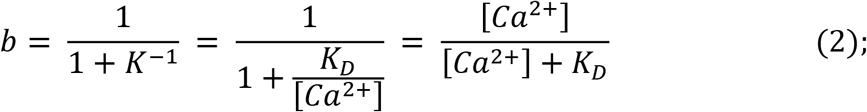

and

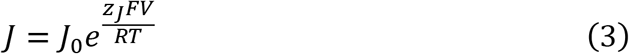

Substituting *b* and *J* into Equation (1), the Ca^2+^-dependent voltage sensor activation for Model I is given by the equation

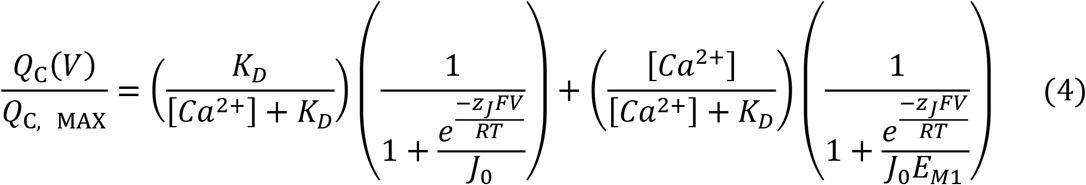

Thus, the *Q*_C_(*V*) curves are determined by the proportion of two functional VSD populations with a distinctive effect (unliganded effect or Ca^2+^-saturated effect) Consequently, the *Q*_C_(*V*) curves are represented by a weighted sum of two Boltzmann functions.

Meanwhile, for the concerted CTD-VSD interaction Scheme II (Model II), the *Q*_C_(*V*, [Ca^2+^]) curves would be determined using the general equation:

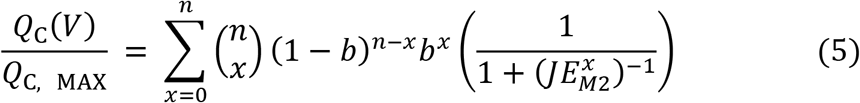

The expression in the first bracket represents the fraction of VSD belonging to a channel with *x* (0 to 4) Ca^2+^ bound, according to a binomial probability distribution. Thus, the *Q*_C_(*V*) curves result in a weighted sum of five distinct Boltzmann functions corresponding to the five possible R-A transitions (***Figure 2—figure supplement 1C***). By stating *n* = 4 because the tetrameric symmetry of the channels, and substituting *b* and *J* into the previous equation (Equation 5) we have

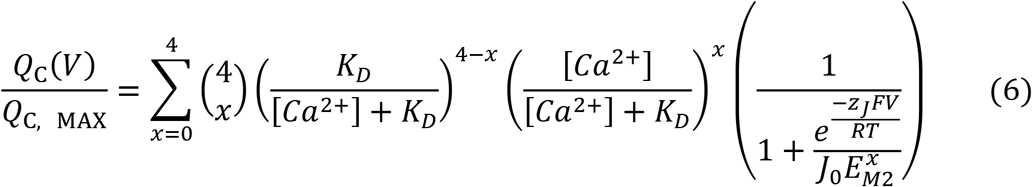

It should be noted that at limiting Ca^2+^ conditions, both schemes become equivalent where the VSD activation is characterized by a single Boltzmann function. At zero Ca^2+^, the *Q*_C_(*V*) curves are described by

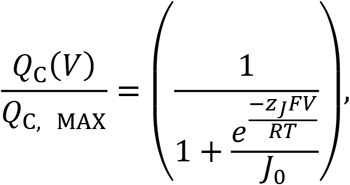

whereas Ca^2+^ saturating concentration *J* is multiply by the allosteric factor *E*, where 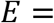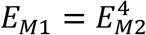 depending on the model (Model I or Model II):

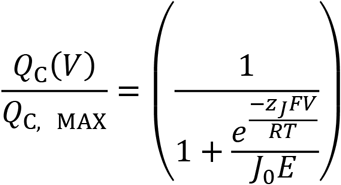

Given that each α-subunit has two Ca^2+^-binding sites, we expanded the CTD-VSD interaction Scheme II (***Figure 2—figure supplement 1C***) considering the existence of two Ca^2+^-binding sites (***Figure 2—figure supplement 1D,E***). The Model II includes the energetic contribution of RCK1 and RCK2 Ca^2+^-sites to the VSD activation. The factor *E* = *E*_*S*1_ * *E*_*S*2_ where *E*_*S*1_ and *E*_*S*1_ are the allosteric coupling between the VSD and the RCK1 Ca^2+^-site and RCK2 Ca^2+^-site, respectively. The *K*_1_ and *K*_2_ constants define the bound/unbound transition for each RCK1 and RCK2 sites being *K*_1_ = [Ca^2+^]/ *K*_*D*1_ and*K*_2_ = [Ca^2+^]/ *K*_*D*2_. Assuming that the Ca^2+^ sensors of distinct α-subunit do not interact, we only consider intrasubunit cooperativity between the RCK1 and RCK2 sites defined by the factor *G*. Thus, the occupancy of one RCK site will affect Ca^2+^-binding equilibrium to the other RCK site in the α-subunit (*GK*_1_ and *GK*_2_) (***Figure 2—figure supplement 1E***). The equilibrium *J* of the VSD increase *E*_*S*1_-fold and *E*_*S*2_-fold for the each Ca^2+^ bound to RCK1 and RCK2 sites, respectively, reaching to 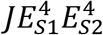 when the eight Ca^2+^ sites are occupied.

## Legends Supplementary Figures

**Figure 1—figure supplement 1. Ca^2+^ increase the slow component of the OFF gating currents.** (**A**) Gating current (*I*_*G*_) recordings evoked by 200 mV pulses with different durations (from 0.2 to 4 ms) at 0 and 10 μM [Ca^2+^] conditions, respectively. (**B-C**) The top panels show the superimposed traces of the *I*_*G*_-OFF recorded at −90 mV evidencing a decrease in amplitude and a slower decay of the OFF current as the duration of the pulse increases at 10 μM Ca^2+^. The dashed line represents the baseline for each experiment. *I*_*G*_-OFF were fitted with an exponential function of two-components (fast and slow components). *I*_*G*_-OFF traces at 0.2 ms and 4 ms pulse duration are displayed for each Ca^2+^ condition. Orange and blue lines correspond to fast and slow components of the two-exponential fits, respectively: (**B**) “zero” Ca^2+^ (*τ*_*F*_ = 10 μs and *τ*_*S*_= 44 μs) and (**C**) 10 μM Ca^2+^ (*τ*_*F*_ = 25 μs and *τ*_*S*_ = 212 μs). (**D-E**) The relative amplitude of the OFF-charge components, the fast (*Q*_*F*_) and slow (*Q*_*S*_) charge components were plotted against the pulse duration and fitted with an exponential function representing the time course of the opening of the channel: (**D**) “zero” Ca^2+^ (*τ*_0Ca_^2+^ = 1.8 ms) and (**E**) 10 μM Ca^2+^ (*τ*_10μM_ = 536 μs).

**Figure 2—figure supplement 1. Kinetic models of the VSD activation according to the CTD-VSD interaction schemes.** (**A**) Sub-scheme describing calcium and voltage allosteric interaction for closed channels. The VSD transit between two resting (R) and active (A) configuration governed by the equilibrium constant *J*, whereas each Ca^2+^ site undergoes unbound (U) - Ca^2+^ bound (B) transitions governed by the equilibrium constant *K*. The allosteric factor *E* accounts for the coupling between the calcium and voltage sensors (CTD-VSD) (**B-C**) VSD kinetic models in presence of Ca^2+^ according to CTD-VSD interaction schemes I and II (***Figure 2A,B***), respectively, where the vertical transitions (R-A) represent the VSD movement and the horizontal transitions (R-R and α-A) are Ca^2+^-binding reactions when the VSD is in the resting or active conformation. For the Scheme I (**B**), each VSD can undergo R_0_-A_0_ or R_1_-A_1_ transitions depending on the unbound or bound state of the Ca^2+^ site in the α-subunit, respectively. Thus, the R_1_-A_1_ equilibrium is defined by *J* increased *E*_*M*1_-fold *JE*_*M*1_. For the Scheme II (**C**), the R_0_-A_0_ to R_4_-A_4_ transitions represent the VSD equilibrium with 0, 1, 2, 3, and 4 occupied Ca^2+^ sites in the channel. Thus, for each Ca^2+^ bound the equilibrium constant *J* increase *E*_*M*2_-fold reaching to 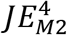 when the four Ca^2+^ sites are occupied. The thickness of the arrows indicates the probability of transitions. (**D**) General sub-scheme of the CTD-VSD interaction including two Ca^2+^ sites for each CTD (RCK1 and RCK2 sites). For each RCK1 and RCK2 site the unbound-Ca^2+^ bound transitions are governed by the equilibrium constants *K*_1_ and *K*_2_. The factor *G* describe the cooperativity between the sites within the same α-subunit; and the *E*_*S*1_ and *E*_*S*2_ factors define the allosteric coupling between the RCK1 and RCK2 sites and the VSD, respectively. (**E**) Schematic representation of VSD kinetic model according to the extended version of the Scheme II (**C**) accounting for both RCK1 and RCK2 Ca^2+^-sites on each α-subunit. For sake of simplicity, are only depicted the VSD transitions depending on the unbound or bound state of the RCK sites within the same α-subunit: RCK1 site (R_1,0_-A_1,0_), RCK2 site (R_0,1_-A_0,1_) and both sites (R_1,1_-A_1,1_).

